# Regulatory effects of beta-2-microglobulin on reactive oxygen species generation in polymorphonuclear cells

**DOI:** 10.1101/2025.10.31.685808

**Authors:** Sofie Espersen Poulsen, Ellen Staudinger, Maria Abildgaard Steffensen, Peter Østrup Jensen, Frederik Vilhardt, Mogens Holst Nissen

## Abstract

Beta-2-microglobulin (β2m) constitutes the invariant light chain of the major histocompatibility complex class I (MHC I) and is present on all nucleated cells, as well as in biological fluids, platelets and neutrophil granules. Immune cells can increase their production and secretion of β2m in response to cytokines such as IFN-γ, IFN-α, and TNF-α. Elevated β2m levels can be detected in various inflammatory, autoimmune, infectious and malignant diseases, often correlating with poor prognosis and reduced survival. So far, no specific functions have been agreed on for soluble β2m.

In the present study, we investigated the effects of exogenous β2m and two proteolytically processed forms, cK58β2m and dK58β2m, on ROS production in polymorphonuclear leukocytes (PMNs). β2m was shown to enhance ROS generation triggered by latex beads, while cK58β2m and dK58β2m suppressed both baseline ROS levels and responses to latex beads, TNF-α, and fMLF. Among these, dK58β2m was generally the most potent inhibitor. The inhibitory effect of cK58β2m and dK58β2m was also confirmed in DMSO-differentiated HL-60 cells, a neutrophil-like cell line. Furthermore, treatment with dK58β2m impaired the recruitment of p47phox and p67phox to the plasma membrane following fMLF stimulation, two key subunits of the NADPH oxidase complex responsible for superoxide anion production in neutrophils. This may explain the reduced ROS production observed in cells treated with dK58β2m.

Taken together, these results suggest that β2m, cK58β2m and dK58β2m can regulate ROS generation in PMNs and highlight a potential role for β2m in modulating the innate immune response through control of ROS production.

## Introduction

Neutrophil granulocytes are the most abundant immune cell in human blood, accounting for about 50-70% of circulating leukocytes. They play a crucial role in acute inflammation and serve a vast array of specialized functions^1–3^. Neutrophils can produce large amounts of reactive oxygens species (ROS) in a process known as the respiratory burst. The production of ROS can be induced by various pro-inflammatory signals including cytokines, bacterial products and pathogen phagocytosis. ROS serve as a key element of the innate immune defense against bacterial and fungal infections and also act as important immunomodulatory signaling molecules^4,5^. ROS can be released both extracellularly into the local environment and intracellularly within phagosomes following phagocytosis. Tight regulation of ROS generation is necessary to prevent excess tissue damage and uncontrolled inflammation.

In neutrophils, ROS production is mediated by the phagocyte NADPH oxidase complex, a multi-component enzyme that catalyzes the NADPH-dependent reduction of oxygen to superoxide anion. In the resting state, the NADPH oxidase subunits are distributed between the cytosol and membrane. Upon activation by agents such as the bacterial chemotactic peptide N-formyl-methionyl-leucyl-phenylalanine (fMLF), the cytosolic components (p47phox, p67phox, p40phox and Rac2) translocate to the membrane and associate with the membrane-associated subunits (NOX2 and p22phox) to form the active enzyme complex^4,5^. Both p47phox and p67phox play an important role in facilitating the recruitment of the other cytosolic components^6^.

Beta-2-microglobulin (β2m) is a small, single-chain protein expressed by all nucleated cells as part of the MHC class I complex^7,8^. Monomeric β2m is found in biological fluids^9^, in platelets^10^ and within neutrophils grenules^11–13^, and can also be secreted by immune cells in response to various stimuli including pro-inflammatory cytokines^14^. Upon activation with agents such as fMLF and PMA, neutrophils release granular β2m^15–17^, although its specific function in this context remains unclear. Elevated serum levels of β2m have been reported in a wide range of conditions, including inflammatory diseases^18–21^, viral infections^22,23^, malignancies^24–26^ and chronic kidney disease^27^. Increased β2m has also been detected in the gingival crevicular fluid (GCF) of patients with periodontitis^28^. In addition, high β2m levels are associated with atherosclerosis and cardiovascular disease, and with increased mortality in these patients^29^. In multiple myeloma, β2m is widely recognized as a key prognostic marker^30,31^.

In 1979, Plesner and Wiik first identified a modified variant of β2m that displayed alpha mobility in immunoelectrophoresis and was present in the serum of patients with systemic lupus erythematosus and rheumatoid arthritis^32^. This variant was later detected in patients with several malignant diseases as well^33–36^. Nissen *et al*. showed that it lacked Lys58 in the disulfide loop and named it desLys58-β2m (dK58β2m)^37^. It is generated by proteolytic cleavage of β2m mediated by C1s complement^38^, followed by removal of Lys58 by a carboxypeptidase B-like activity^39^. The C1s cleaved intermediate is referred to as Lys58-cleaved β2m (cK58β2m). Biophysical characterization revelated that dK58β2m adopts a conformation with amyloidogenic features^40–42^. Compared with native β2m, both dK58β2m and cK58β2m exhibit increased heterogeneity and reduced stability under physiological conditions^43^. Importantly, high levels of cleaved β2m in blood have been found to correlate with disease activity in autoimmune^32^ and malignant disorders ^33,34,36,44^. In patients with end-stage renal diseases, dK58β2m has also been detected in serum and was found to be associated with the use of less biocompatible dialysis membranes^45^.

Only a few studies have explored the biological significance of dK58β2m^46–48^. Exogenous dK58β2m has been shown to augment the generation of specific cytotoxic activity in a one-way mixed lymphocyte culture by promoting an early increase in IL-2 release^48^. In combination with interferon-γ (IFN-γ), dK58β2m has been reported to induce nitric oxide production and trigger pro-apoptotic effects in myeloid cell lines^46,49^. Co-stimulation with IFN-γ and either β2m or dK58β2m also increases mitochondrial reactive oxygen species (mtROS) production^46^. Moreover, dK58β2m together with IFN-γ can activate the cyclic GMP–AMP synthase (cGAS)–stimulator of interferon genes (STING) pathway^46^.

In this study, we show that cK58β2m and dK58β2m inhibit ROS production in human polymorphonuclear leukocytes (PMNs, predominantly neutrophils) and DMSO-differentiated HL-60 cells. The reduction in ROS generation could not be explained by these proteins acting as direct superoxide scavengers. Instead, dK58β2m was found to impair assembly of the NADPH oxidase at the membrane, demonstrated by decreased translocation of p47phox and p67phox following fMLF stimulation. Furthermore, the native form of β2m was shown to enhance latex bead-induced ROS production in PMNs but did not show a similar stimulatory effect on ROS generation triggered by soluble single stimuli.

## Materials and methods

### Protein expression, cleavage and purification

Human recombinant β2m was produced in the ExpiCHO^™^ expression system (Thermo Fisher Scientific, Gibco^™^) cK58β2m and dK58β2m were generated by cleavage of β2m, and all variants were purified as described previously with minor modifications^46,50^. The proteins were first separated on a G75 Sephadex gel filtration column. The cK58β2m variant was then isolated by chromatofocusing on a Polybuffer Exchanger 94, while dK58β2m was purified by ion exchange chromatography on a Resource Q column. Finally, the samples were up-concentrated by ammonium sulphate precipitation and subsequently desalted by passing the solution through PD-10 desalting columns.

### Cells and reagents

The human promyeloblast cell line HL-60 (CCL-240^™^, ATCC®) was cultured in RPMI 1640 supplemented with GlutaMAX^™^ with 1% penicillin-streptomycin and 10% fetal bovine serum (FBS; A5256701, Thermo Fisher Scientific, Gibco^™^). HL-60 cells were set to differentiate in complete medium with 1.3% DMSO for 5 days at 37°C with 5%Co_2_^51^. Differentiation was verified based on morphological changes, NOX2 expression and phagocytic activity (data not shown).

Dihydrorhodamine 123 (DHR-123), fMLF, superoxide dismutase (SOD), diphenylene iodonium (DPI), luminol, 3 µm polystyrene latex beads, horseradish peroxidase type II (HRP II), xanthine oxidase, and hypoxanthine were all purchased from Sigma-Aldrich. Human recombinant tumor necrosis factor alpha (TNF-α) was purchased from R&D systems, and WST.1 was purchased from Santa Cruz Biotechnology. Rabbit anti-p67phox (#PA537323), goat anti-GADPH-HRP (#PA1087), rabbit anti-Na/K-ATPase subunit alpha-1 (#MA532184), and goat anti-rabbit-HRP (#31460) were all purchased from Invivogen^™^, Thermo Fisher Scientific. The rabbit anti-p47phox (‘B’) antibody was kindly provided by Rikard Holmdahl, Karolinska Institute.

### Isolation of human peripheral blood PMNs

Human PMNs were isolated from heparinized whole-blood samples collected from healthy donors as described previously^52^. Donors were kept completely anonymous, with samples being marked only for age and gender in accordance with Danish legislation and ethical guidelines, and neither samples nor cells were stored.

### Flow cytometric detection of ROS production

Intracellular ROS in PMNs were detected as oxidation of DHR-123 by flow cytometry. Cells were incubated with 50 µg/ml β2m, cK58β2m or dK58β2m either alone or in combination with 10 ng/ml TNF-α or 1 µM fMLF, and 40 µM DHR-123 for 1 h at 37°C with 5% CO_2_. As a control, cells were treated with 100 nM DPI. The cells were run on a CytoFLEX S (Beckman Coulter) and data was analyzed using FlowJo (v10.10.0).

### Luminol-amplified chemiluminescent ROS detection

Total ROS production in PMNs and DMSO differentiated HL-60 cells (dHL-60) was analyzed by luminol-amplified chemiluminescence. In one setup, ROS production by PMNs in response to treatment with 3 µm polystyrene latex beads was studied. Cells were treated with 50 µg/ml β2m, cK58β2m or dK58β2m in combination with 70 µM luminol and latex beads in a ratio of 5:1 or 3.3:1 relative to the cell number. Chemiluminescence was detected every 20 s on a Wallac Victor2 1420 plate reader (Perkin Elmer) for a total of 1 h at 37°C.

In another setup, fMLF-induced ROS production by PMNs or dHL-60 cells was studied. Cells were pre-incubated with β2m, cK58β2m or dK58β2m at different concentrations together with 50 µM luminol for 5 min at 37°C. For HL-60 cells, HRP II (2.5 U/ml) was added together with luminol to ensure efficient luminol oxidation as they show lower peroxidase levels than PMNs^53^. Treatment with 50 U/ml SOD was used as a negative control. Chemiluminescence was detected every 6 s for 3 min (PMNs) or 2.5 min (dHL-60) on a FlexStation 3 (Molecular Devices) at 37°C. After 20 s, 10 µM fMLF was added to the wells.

### Xanthine oxidase/hypoxanthine system

Superoxide anion scavenging effects of β2m, cK58β2m and dK58β2m were investigated using the cell-free xanthine oxidase/hypoxanthine system^54^. Each β2m variant (50 µg/ml) was incubated with 0.25 mM hypoxanthine, 3 mU/ml xanthine oxidase and 500 µM WST.1 for 20 min at 37°C. SOD (50 U/ml) was used as positive control. Absorbance at 450 nm was recorded every 30 s using a FlexStation 3 microplate reader (Molecular Devices).

### Western blot

DMSO-differentiated HL-60 cells (5×10^6^) were pre-incubated with 50 µg/ml β2m or dK58β2m for 5 min, followed by stimulation with 1 µM fMLF for 1 min at 37°C. Cytosolic and membrane-associated proteins were isolated using the Mem-PER™ Plus Membrane Protein Extraction Kit (Thermo Fisher Scientific) according to the manufacture’s recommendations. Halt™ Protease and Phosphatase Inhibitor Cocktail (Thermo Fisher Scientific) was added at all membrane and cytosol extraction steps. Protein concentrations were determined using Pierce^™^ BCA Protein Assay (Thermo Fisher Scientific), and equal amounts of protein (5 µg protein/well) were separated on 10% SDS-PAGE gels and transferred onto PDVF membranes. Membranes were blocked in 2% BSA prepared in Tris-buffered saline with 0.1% Tween 20 (TBS-T) and then incubated overnight at 4°C with the following primary antibodies: anti-p47phox (1:2000), anti-p67phox (1:1000), anti-GAPDH-HRP (1:4000), and anti-Na/K-ATPase subunit alpha-1 (1:20,000). After washing with TBS-T, membranes were incubated with HRP-conjugated goat anti-rabbit secondary antibody (1:20,000) for 1 h at room temperature, except for GAPDH detection, as the primary antibody was already HRP-conjugated. Protein bands were developed using Amersham ECL Prime Chemiluminescence Reagents (Cytiva) and imaged on a ChemiDoc^™^ MP Imaging System (Bio-Rad). Band intensities were quantified using ImageJ (Fiji) software (version 1.54p). Protein levels were normalized to β-actin in cytosolic fractions and to Na/K-ATPase subunit alpha-1 in membrane fractions. Data are presented as fold change in densitometric signal, expressed as the relative membrane-to-cytosol ratio for each protein.

### Statistical analysis

Statistical analysis was performed using GraphPad Prism 10. Normality tests (D’Agostino and Pearson test or Shapiro-Wilk test) were run to determine the appropriate statistical test. Comparisons were made using paired student’s t test, repeated-measures ANOVA, one-way ANOVA or Kruskal-Wallis testfollowed by Dunnett’s or Dunn’s multiple comparison test as applicable. A p-value <0.05 was considered statistically significant.

## Results

### β2m enhances, whereas the cleaved variants, cK58β2m or dK58β2m, inhibit latex bead-induced ROS production in PMNs

During neutrophil degranulation, β2m is released into the extracellular space^15–17^. The functional significance of this ‘free’ β2m during inflammatory responses remains unclear. To investigate whether β2m influences neutrophil effector functions, we examined the effects of native β2m and its cleaved variants, cK58β2m and dK58β2m, on ROS production in human PMNs.

Phagocytosis by neutrophils is closely linked to their generation of ROS^5,55^. To investigate, if the β2m variants modulate ROS production triggered by phagocytosis, we stimulated freshly isolated PMNs with 3 µm latex beads either alone or combined with 50 µg/ml of each β2m variant. ROS production was followed for 1 hour using luminol-enhanced chemiluminescence. We found that native β2m increased ROS production to 133 ± 26% compared to beads alone (Figure 1). In contrast, the cleaved variants were shown to suppress the ROS response, with cK58β2m reducing it to 57 ± 32% and dK58β2m to 74 ± 29% (Figure 1).

**Figure 1.**
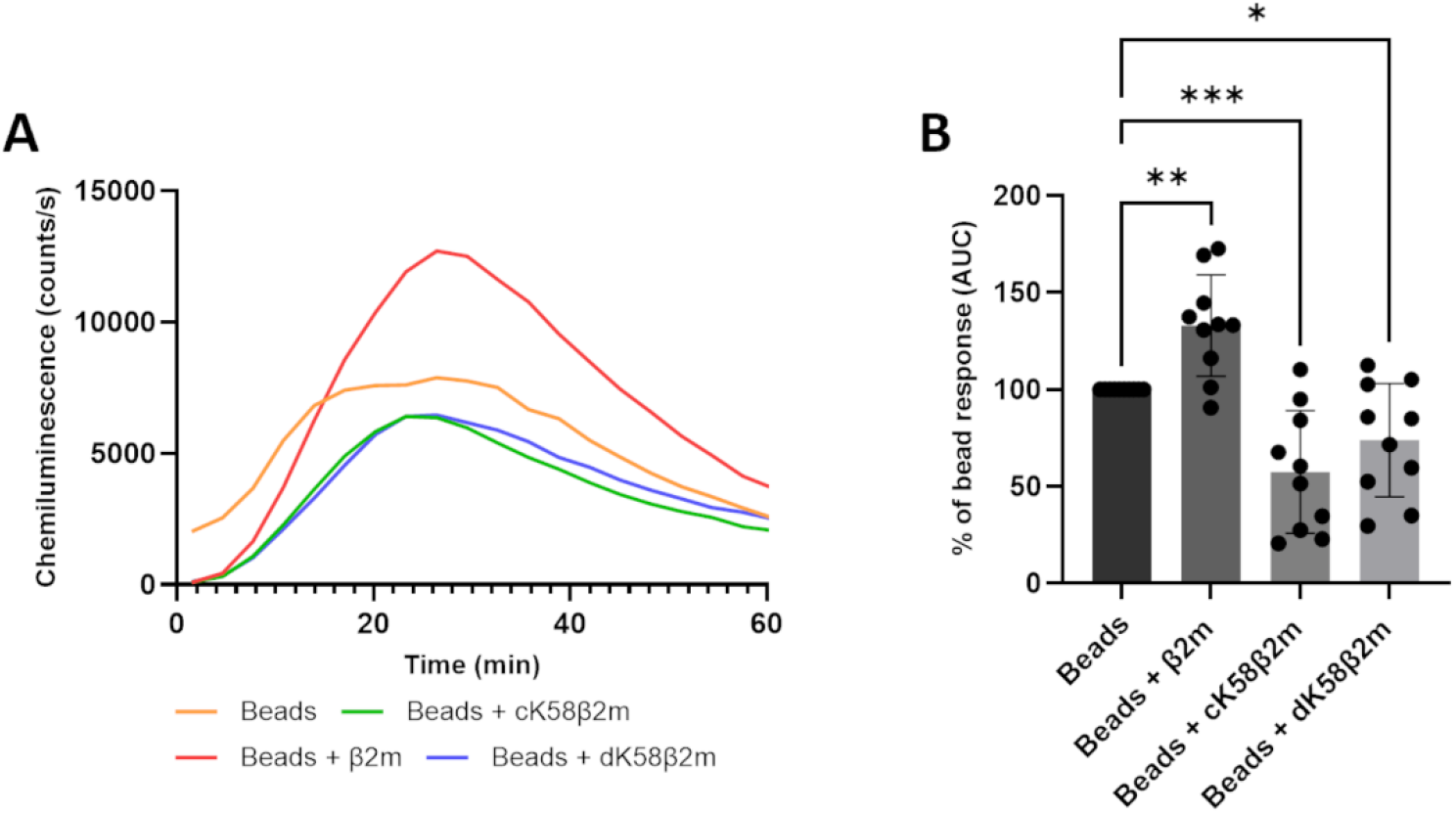
β2m increases, while cK58β2m and dK58β2m inhibit bead-induced ROS response in PMNs. PMNs were treated with 50 µg/ml β2m, cK58β2m or dK58β2m in combination with latex beads (5:1 or 1:3.3 latex bead to PMN ratio). ROS production was followed for 1 h by luminol-enhanced chemiluminescence. (A) Representative plot of the ROS response. (B) Results are expressed as mean ± SD, as relative AUC. n=10 donors, *P<0.05, **P<0.01, ***P<0.001. Group comparisons were analyzed by repeated-measures one-way ANOVA and Dunnett’s post hoc test.

### cK58β2m and dK58β2m inhibit ROS production in PMNs

Neutrophil ROS production can be triggered by a wide range of stimuli. To gain a deeper understanding of how β2m affects ROS generation in these cells, we investigated its influence on both baseline ROS levels and ROS production in response to two known stimulants: the bacterial chemoattractant peptide fMLF and the pro-inflammatory cytokine TNF-α^56^.

Intracellular ROS levels in freshly isolated PMNs were measured via flow cytometry as DHR-123 oxidation. We found that both cleaved β2m variants significantly suppressed baseline ROS production (Figure 2A), reducing it to 53 ± 20% for cK58 β2m and 64 ± 30% for dK58β2m relative to the untreated control. For the native protein, no significant difference was observed. Interestingly, both cK58β2m and dK58β2m were shown to reduce baseline ROS to a level similar to DPI, a known NADPH oxidase inhibitor (p=0.9458 and p=0.4022, respectively) (Figure 2A).

**Figure 2.**
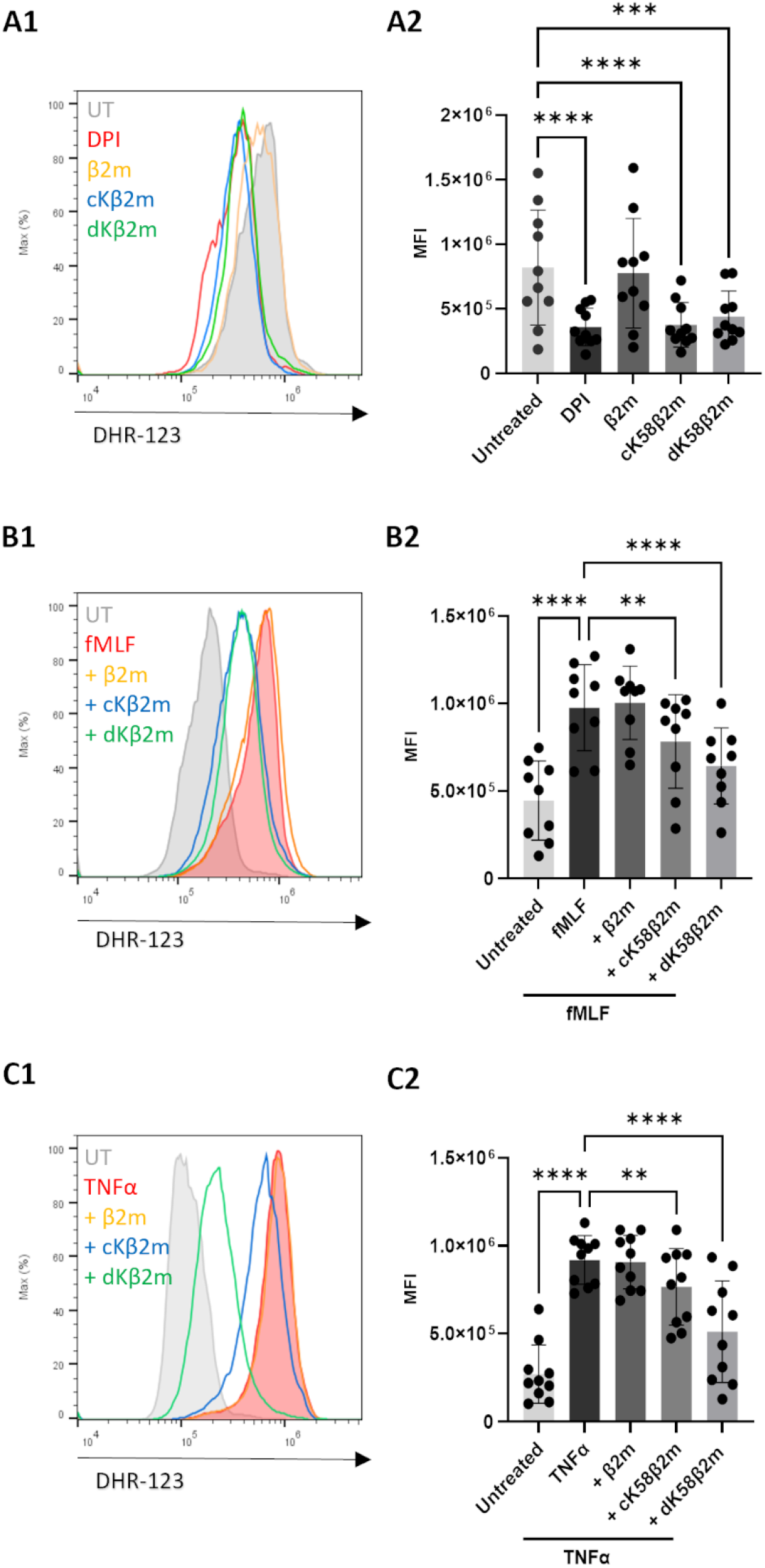
cK58β2m and dK58β2m inhibit intracellular ROS production in PMNs. PMNs were treated with 50 µg/ml β2m, cK58β2m or dK58β2m either alone (A) or in combination with 1 µM fMLF (B) or 10 ng/ml TNF-α (C) for 1 h. DPI (200 nM, NADPH oxidase inhibitor) treatment was used as a control. Intercellular ROS levels were detected using DHR-123 (40 µM) by flow cytometry. (A1-C1) Representative histograms of the DHR-123 signal. (A2-C2) Results are expressed as mean MFI ± SD. n=10 (A2 and C2) and n=9 donors (B2), . **P<0.01, ***P<0.001, ****P<0.0001. Group comparisons were analyzed by repeated-measures one-way ANOVA and Dunnett’s post hoc test.

When looking at stimuli-induced ROS responses, cK58β2m and dK58β2m were found to inhibit both fMLF- and TNF-α-induced ROS production, while the native form had no effect (Figure 2B and C). Notably, dK58β2m exerted a stronger inhibitory effect than cK58β2m in both cases: cK58β2m reduced fMLF- and TNF-α-induced ROS production to 78 ± 15% and 82 ± 13%, respectively, whereas dK58β2m lowered these responses to 66 ± 17% and 52 ± 24%. The inhibitory effect of cK58β2m and dK58β2m on TNF-α-induced ROS production was found to be sex-dependent, with PMNs from female donors exhibiting a significantly greater reduction in ROS levels compared to those from male donors (Figure S2).

The inhibitory effect of the cleaved β2m forms on fMLF-induced ROS production were further validated using luminol, a probe that detects both intracellular and extracellular ROS (Figure 3B and C). Using this probe, dK58β2m was shown to suppress ROS production in a dose-dependent manner, achieving a maximum reduction of 53 ± 11% at a concentration of 50 µg/ml. Interestingly, unlike the DHR-123 findings, 50 µg/ml β2m was found to decrease the fMLF-induced ROS response when measured with luminol (to 86 ± 10%) (Figure 3A). Pre-treatment with SOD reduced the response to 23 ± 0.5%, indicating that superoxide anion produced by the NADPH oxidase is a major contributor to the observed ROS signal (Figure 3D).

**Figure 3.**
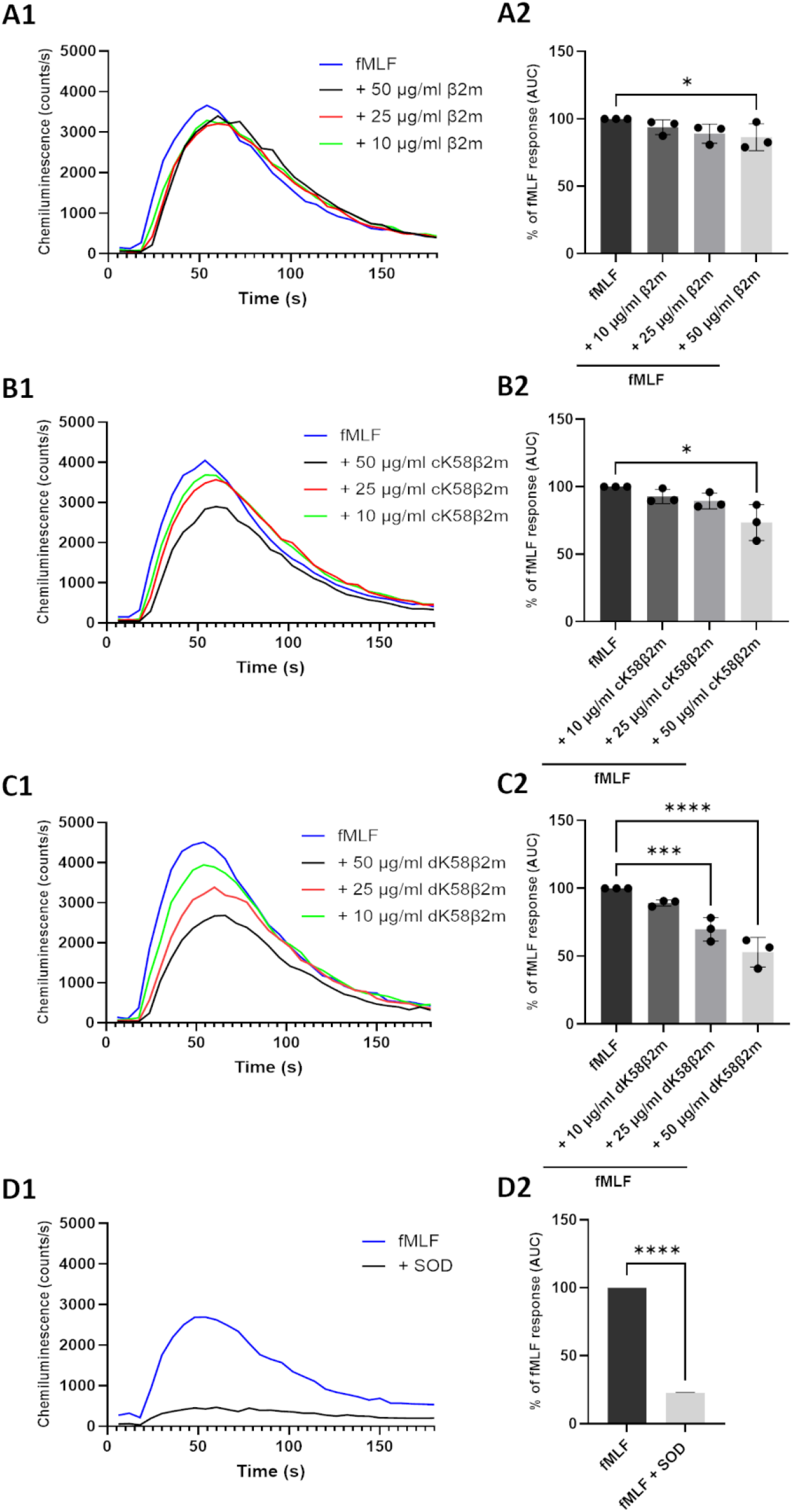
cK58β2m and dK58β2m inhibit fMLF-induced ROS production in PMNs in a dose-dependent manner. PMNs were pre-treated with β2m (A), cK58β2m (B) or dK58β2m (C) for 5 min followed by stimulation with 10 µM fMLF. Pre-treatment with 50 U/ml SOD was used as control (D). fMLF-induced ROS production was followed over a 3-min period by luminol-enhanced chemiluminescence. (A1-D1) Representative plots of the ROS response. (A2-D2) Results are expressed as mean ± SD, as percentage of the fMLF-response. n=3 donors. **P<0.01, ***P<0.001, ****P<0.0001. Comparisons between groups were analyzed with repeated-measures one-way ANOVA and Dunnett’s post hoc test (A-C) or paired t test (D).

Taken together, these experiments demonstrate that both cK58β2m and dK58β2m inhibit ROS production in PMNs independent on the type of stimulation or the detection probe used. In contrast, 50 µg/ml β2m exhibits only a slight inhibitory effect on fMLF-induced ROS production when measured with luminol, and not with DHR-123.

### cK58β2m and dK58β2m do not function as direct ROS scavengers

Since both cK58β2m and dK58β2m were found to suppress ROS production in PMNs, we next thought to investigate whether this effect could be explained by a direct ROS scavenging effect of the proteins. To evaluate this, we used the hypoxanthine/xanthine oxidase system to generate superoxide anion in a cell-free environment. None of the three β2m variants demonstrated significant superoxide-scavenging activity (Figure 4), indicating that the observed inhibitory effect of cK58β2m and dK58β2m on ROS production is likely due to modulation of the cellular ROS response rather than a direct antioxidant effect.

**Figure 4.**
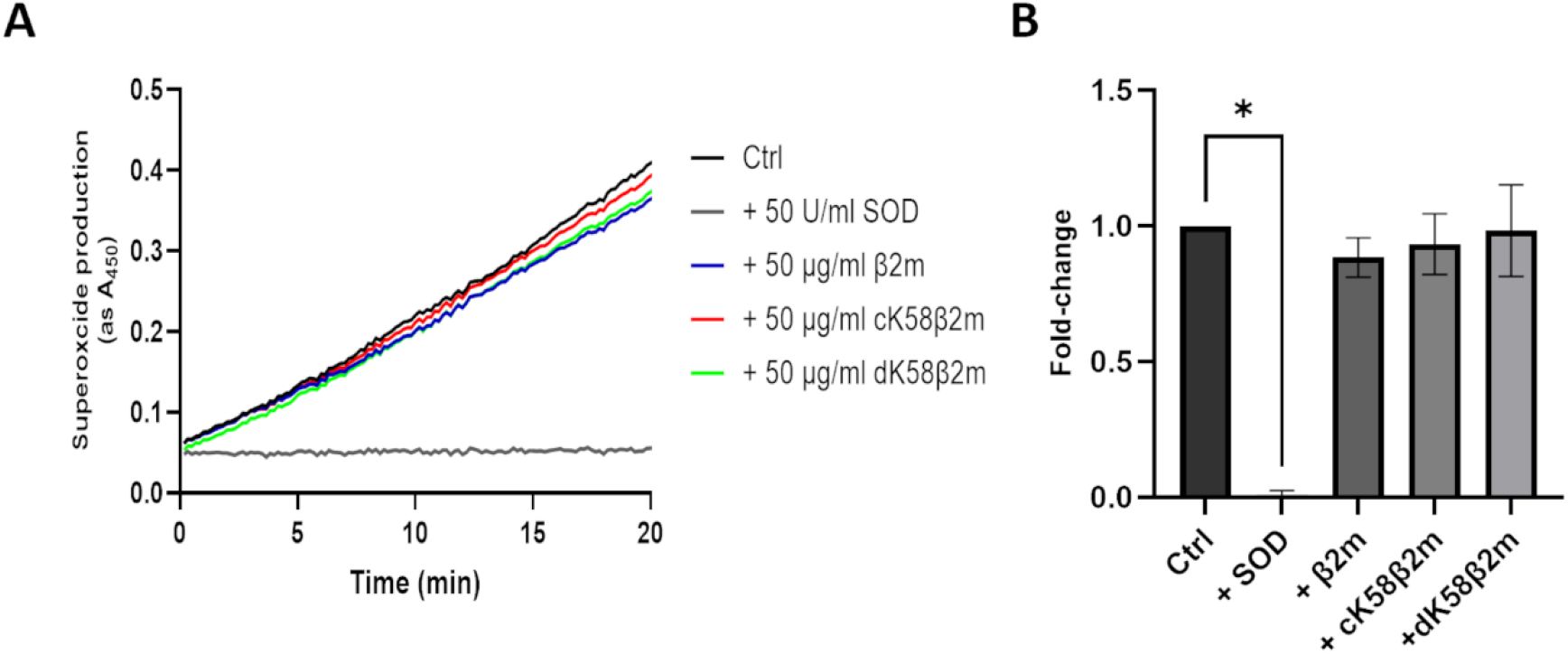
cK58β2m and dK58β2m do not function as direct superoxide scavengers. Superoxide anion was produced in the cell-free xanthine oxidase/hypoxanthine system. The production of superoxide was followed over a 20-min period as chromogenic reduction of WST-1 with 50 U/ml SOD or 50 µg/ml β2m, cK58β2m or dK58β2m. (A) Representative plot of the superoxide anion production. (B) Results are expressed as mean slope ± SD, as fold change of the positive control. n=3, *P<0.05. Comparisons between groups were analyzed with Kruskal-Wallis test and Dunn’s post hoc test.

### cK58β2m and dK58β2m suppress ROS production in the neutrophil-like HL-60 cell-line

To investigate if the cleaved β2m forms exert similar inhibitory effects in a cell line, DMSO-differentiated HL-60 cells (dHL-60) were used as neutrophil-like cells. These cells have no specific granules and therefore the full NADPH oxidase complex is present only on the plasma membrane^57^. dHL-60 cells were pre-treated with 10, 25 or 50 µg/ml of each β2m variant for 5 minutes, after which fMLF-induced ROS production was followed over a 2.5-minute period. Consistent with findings in PMNs, 50 µg/ml of both cK58β2m and dK58β2m inhibited ROS production (Figure 5B and C). Interestingly, a dose-dependent inhibition was apparent for both cleaved β2m variants in dHL-60 cells. Specifically, cK58β2m at 25 and 50 µg/ml significantly reduced the fMLF-induced ROS response to 91 ± 5.7% and 71 ± 6.1%, respectively. Similarly, dK58β2m proved to be the more potent inhibitor in this model, decreasing ROS production to 81 ± 9.6% at 25 µg/ml and 59 ± 9.7% at 50 µg/ml. In dHL-60 cells, treatment with 50 µg/ml β2m also resulted in a statistically significant reduction of the fMLF response (to 87 ± 4.5%) (Figure 5A). Moreover, pre-incubation with SOD nearly abolished the ROS response (reduced to 7 ± 2.5%), confirming that superoxide anion was the main contributor to the detected signal (Figure 5D). Thus, dHL-60 cells seem to be a suitable model to study the effects of cK58β2m and dK58β2m on ROS production in further detail.

**Figure 5.**
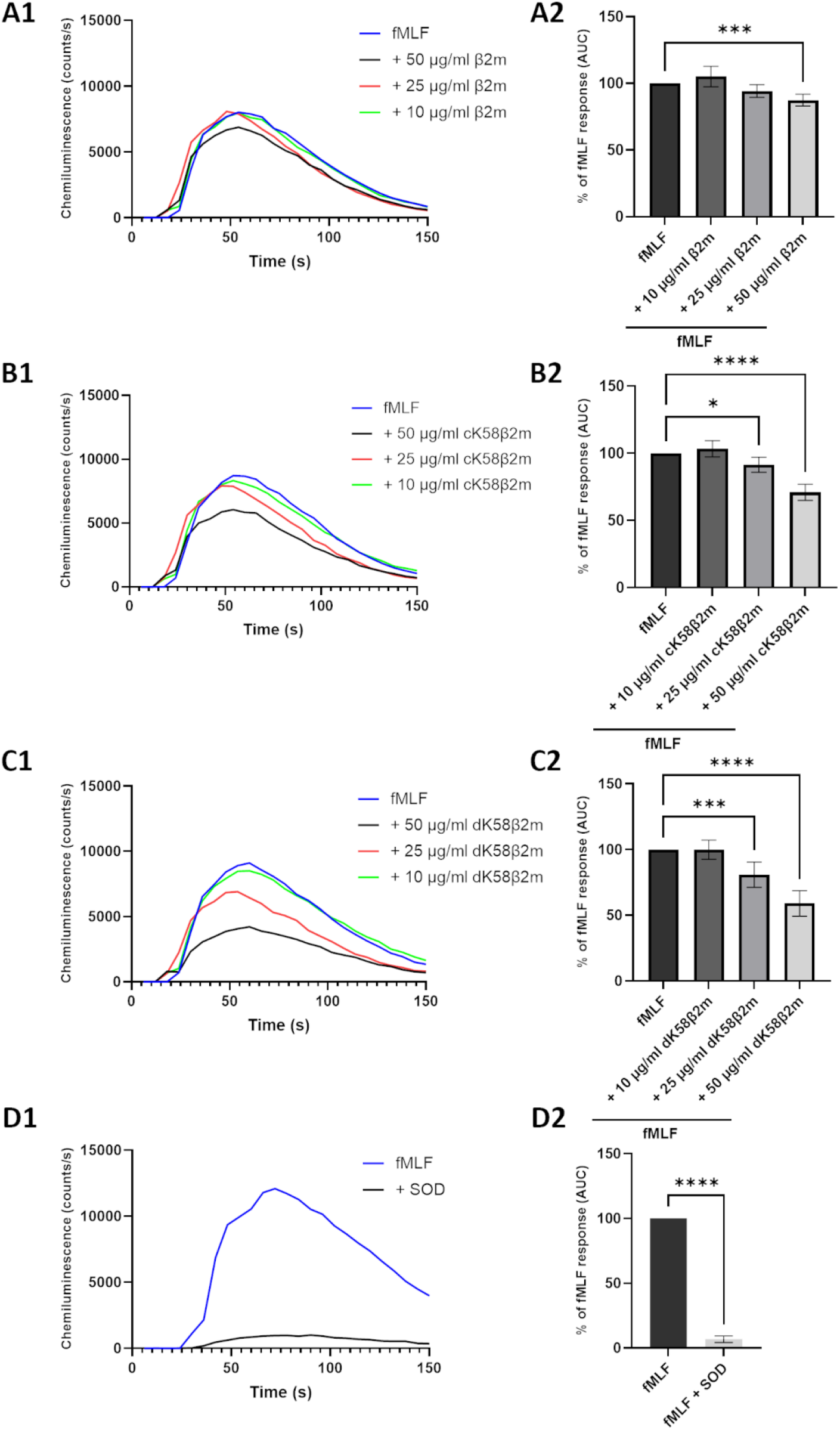
cK58β2m and dK58β2m inhibit fMLF-induced ROS production in dHL-60 cells. dHL-60 cells were pre-treated with β2m (A), cK58β2m (B) or dK58β2m (C) at different concentrations for 5 min followed by stimulation with 10 µM fMLF. Pre-treatment with 50 U/ml SOD was used as negative control (D). fMLF-induced ROS production was followed over a 2.5-min period by luminol-enhanced chemiluminescence. (A1-D1) Representative plots of the ROS response. (A2-D2) Results are expressed as mean ± SD, as percentage of the fMLF-response. n=6, *P<0.05, ***P<0.001, ****P<0.0001. Comparisons between groups were analyzed with one-way ANOVA and Dunnett’s post hoc test (A2-C2) or paired t test (D2).

### dK58β2m inhibits ROS production by impairing assembly of the phagocyte NADPH oxidase complex in dHL-60 cells

The phagocyte NADPH oxidase is an essential enzyme for neutrophil respiratory burst, responsible for generating superoxide anion. It is composed of six subunits, two membrane-bound and four soluble subunits, that assemble at the membrane upon activation^58,59^. To investigate whether the inhibitory effect of dK58β2m on ROS production may result from altered NADPH oxidase assembly and function, we studied the translocation of key subunits, p47phox and p67phox, from the cytosol to the membrane. Cytosolic and membrane protein fractions were collected from dHL-60 pre-treated with 50 µg/ml β2m or dK58β2m for 5 minutes followed by stimulation with fMLF. We found that pre-treatment with dK58β2m significantly reduced both p47phox (0.5 ± 0.26-fold) and p67phox (0.41 ± 0.06-fold) translocation to the membrane following fMLF stimulation (Figure 6). In contrast, β2m treatment decreased p67phox translocation (0.68 ± 0.06-fold), whereas a tendency to more p47phox in the membrane fraction was observed (1.4 ± 0.07-fold, p=0.0548) (Figure 6). Taken together, these findings indicate that dK58β2m impairs the recruitment of both p47phox and p67phox subunits to the membrane, which may explain its inhibitory effect on ROS production in dHL-60 cells.

**Figure 6.**
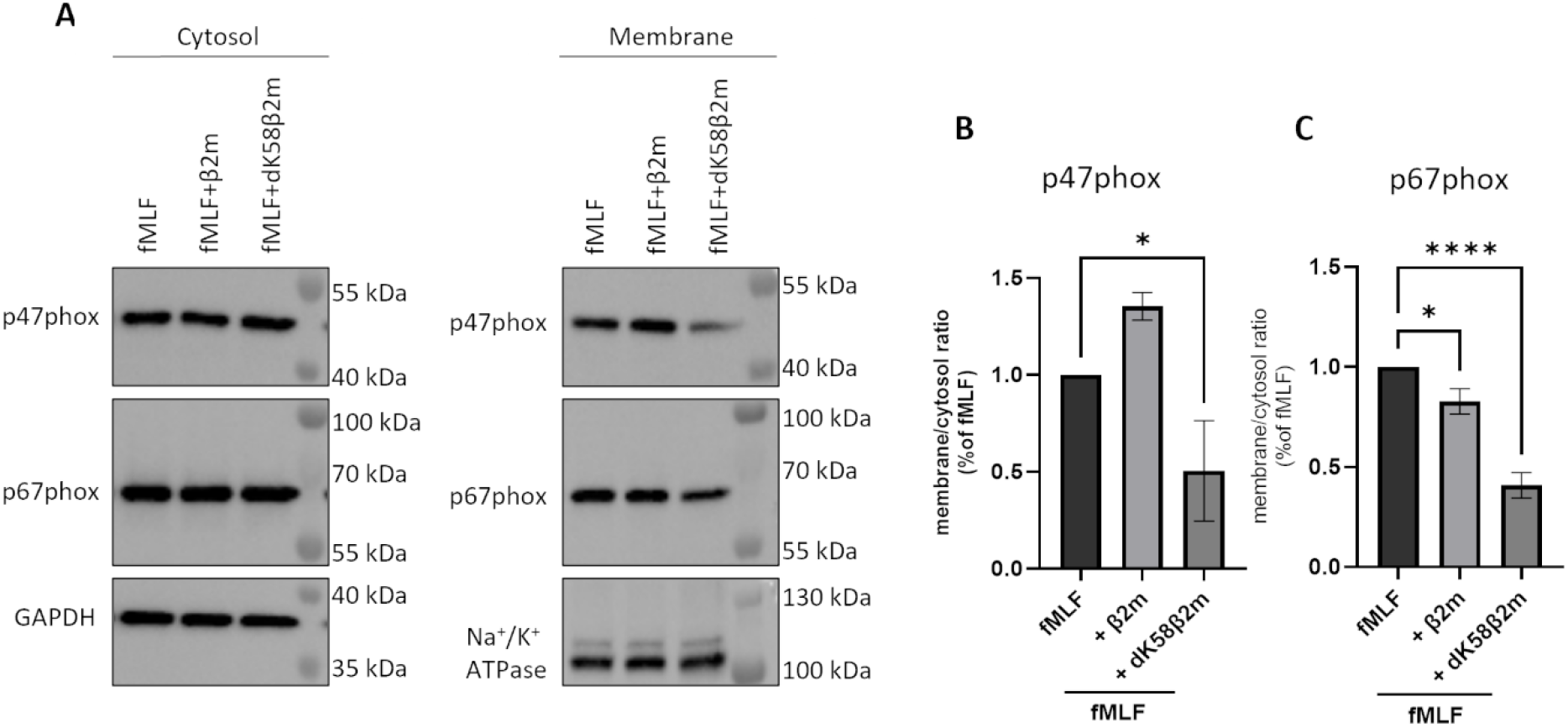
dK58β2m impairs translocation of p47phox and p67phox to the membrane upon stimulation with fMLF in dHL-60-cells. dHL-60 cells were pre-treated with 50 µg/ml β2m or dK58β2m for 5 min followed by stimulation with 1 µM fMLF for 1 min. Cytosolic and membrane protein fractions were collected, and p47phox and p67phox levels were evaluated in each fraction by western blot. (A) Representative blots. (B-C) Relative membrane/cytosol ratio of p47phox (B) and p67phox (C). Results are expressed as mean ± SD, as fold change of the fMLF response. n=3, *P<0.05, ****P<0.0001. Comparisons between groups were analyzed with one-way ANOVA and Dunnett’s post hoc test.

## Discussion

In the present study, we demonstrate that exogenous β2m regulates ROS production in PMNs. While native β2m enhanced ROS generation induced by latex beads, the cleaved forms, cK58β2m and dK58β2m, generally suppressed ROS production. Based on these findings, we propose that β2m may function in an autocrine and/or paracrine manner following neutrophil degranulation, modulating neutrophil effector responses similarly to other granule-derived proteins^60^.

We demonstrated that stimulation of PMNs with exogenous β2m enhanced their ROS response to latex beads. The ability of neutrophils to produce ROS following phagocytosis of latex beads is well established^61,62^. To further explore the effects of β2m, we also examined baseline ROS levels and ROS responses to soluble stimuli such as TNF-α and fMLF. Interestingly, contrary to bead-induced ROS, β2m (at 50 µg/ml) inhibited fMLF-triggered ROS production in both PMNs and dHL-60 cells when measured as luminol oxidation, although this inhibition was not detected using the intracellular ROS probe DHR-123. This observed inhibitory effect may be due to fMLF stimulation preferentially targeting the surface-resident NADPH oxidase, with minimal activation of NADPH oxidase in internal compartments^63^. Examination of NADPH oxidase assembly following fMLF stimulation in combination with β2m showed contrasting results: decreased membrane translocation of p67phox but not of p47phox. The stimulatory effect seen with β2m in combination with latex beads may reflect a requirement for β2m internalization by neutrophils. Alternatively, β2m may enhance phagocytic activity, thereby indirectly increasing ROS production. Finally, it could be speculated that p40phox is activated by latex beads, and acts as an alternative organizer to p47phox, as p40phox does in relation to Fcγ receptor signaling^6,64,65^.

The cleaved β2m forms both exerted an inhibitory effect on ROS production in PMNs. cK58β2m and dK58β2m suppressed both baseline ROS levels and ROS responses induced by latex beads as well as TNF-α and fMLF, indicating that their inhibitory action is not stimulus specific. Results obtained on dHL-60 stimulated with fMLF and the different β2m forms supported our findings in PMNs. In both cell types, dK58β2m was shown to be the most potent inhibitor of ROS formation triggered by fMLF. Since SOD treatment nearly abolished the fMLF-induced ROS response in both PMNs and dHL-60 cells, this strongly suggests that superoxide anion is a major contributor to the detected ROS response. To explore this further, we examined assembly and activation of the phagocyte NADPH oxidase complex in dHL-60 cells. We found that dK58β2m inhibits assembly and thus activation of this complex, which may explain the reduced ROS levels in cells treated with dK58β2m. Tests for direct antioxidant activity of cK58β2m and dK58β2m found no evidence that they scavenge superoxide anion directly.

Bjerrum *et al*. have also studied the effect of β2m variants on ROS production in neutrophils and found no effect of β2m or dK58β2m on PMA-induced superoxide anion production^15^. PMA is considered a strong stimulator of ROS production and in our experiments, it induced ROS responses at levels several-fold higher than those triggered by latex beads (data not shown). Thus, it is likely that PMA activates ROS production at levels that may overwhelm the modulatory effects of β2m, in particular that strong activation of p47phox or p40phox by protein kinase C (PKC) phosphorylation (activated by PMA)^66^ cannot be relieved by the cleaved β2m variants.

Our results indicate that the regulatory effects of β2m on ROS production in PMNs depend on both its concentration and the presence of additional stimuli, consistent with previous findings^10,46,47,67^. The highest β2m concentration tested in our study (50 mg/L) greatly exceeds typical serum levels in healthy individuals (<2 mg/L)^68,69^ as well as in patients with malignant or inflammatory conditions (<10 mg/L)^69^. However, β2m levels can locally reach up to 100 mg/L in fluids such as crevicular fluid, salivary and tear fluid during inflammation^28,70–72^. Therefore, it is possible that β2m concentrations in intercellular fluids and confined spaces can exceed serum levels. Interestingly, increased β2m levels have been reported in the gingival crevicular fluid (GCF), the fluid located in the narrow sulcus between the tooth and gum, of patients with severe periodontitis^28,72^. Periodontitis is a serious bacterial infection of the supporting structures of the teeth^73^. In healthy individuals, β2m concentrations in GCF are typically around 10 mg/L, whereas levels can rise to 100 mg/L in severe cases of periodontitis^28^. Notably, neutrophils are the most abundant leukocyte in the gingival sulcus and periodontal pockets in both healthy and diseased states^74,75^, making it tempting to speculate that they may be one of the main contributors to the elevated β2m levels in GCF observed in patients with periodontitis.

In this context it can be noticed that the level of β2m in GCF is found to correlate with the severity of periodontitis^28^. Our findings suggest that native β2m and its cleaved forms have distinct regulatory effects on ROS production: native β2m enhances latex bead-induced ROS production, whereas the cleaved forms suppress it. It could be speculated that the balance between these β2m forms could affect the clinical course of periodontitis. In support of this hypothesis are reports showing that gain-of-function mutations in C1 complement, which is associated with periodontal Ehlers-Danlos syndrome, lead to more severe and early onset periodontitis^76,77^. In this situation, extracellularly secreted C1r/C1s complement may drive proteolysis of β2m, thereby modulating the inflammatory response.

Of the two cleaved variants, only dK58β2m has been identified in humans, more particular in patients with autoimmune diseases, malignancies, and renal disorders^33–36,45^. Since the level of both β2m and C1s complement component increase during inflammation, we hypothesize that dK58β2m may be present at inflammatory sites, although this remains to be confirmed. Notably, neutrophils themselves have been shown to be incapable of converting β2m into dK58β2m^16^, but human β2m can get cleaved when added to a murine one-way mixed lymphocyte culture^16,48^. It would be interesting to investigate how the combination of β2m with cK58β2m or dK58β2m influences bead-induced ROS production to determine which β2m variant exerts the dominant effect.

Recently, we reported that IFN-γ combined with β2m, and especially dK58β2m, induce increased mtROS and nitric oxide production in macrophage-like cells^46^. This aligns with other studies showing that β2m can promote mtROS generation^78,79^. While this connects β2m and dK58β2m to the production of reactive oxygen and nitrogen species, the mechanisms and functions of mtROS differ from those of NOX2-dependent ROS.

From a general perspective, the proteolysis of β2m at Lys58 may have a broader implication, pointing towards that immunoglobulin domains can serve additional biological functions following proteolytically processing. As a member of the immunoglobulin superfamily, β2m shares the characteristic immunoglobulin fold, and its cleavage by C1s complement component occurs within the D-E loop between the disulfide-linked cysteine residues at position 25 and 80^39,80^. This is partly comparable to the proteolytic generation of the tetrapeptide Tuftsin, derived from residues 289–292 within the C-D loop of the CH2 domain of IgG^81^. Tuftsin is known to exert different biological functions including stimulation of phagocytosis and enhancement of ROS production^82,83^.

Taken together, our findings suggest that β2m released from PMNs during degranulation can enhance bead-induced ROS production, although the exact molecular mechanism remains unclear. In contrast, dK58β2m, a cleaved form of β2m that we speculate is present at inflammatory sites, inhibits assembly and activation of the phagocyte NADPH oxidase, resulting in decreased ROS production by PMNs. This potential auto-regulatory role of cleaved β2m could serve as a feedback mechanism to prevent excessive ROS production and minimize tissue damage during inflammation.

