## Supplemental information for "Regulatory effects of beta-2-microglobulin on reactive oxygen species generation in polymorphonuclear cells"

**Table S1 – β2m increases, while cK58β2m and dK58β2m inhibit bead-induced ROS response in PMNs.** Relative AUC values from individual donors from the experiment shown in figure 1.

| | Beads | Beads + β2m | Beads + cK58β2m | Beads + dK58β2m |
|---|---|---|---|---|
| Donor 1 | 100 | 133.2 | 60.5 | 59.6 |
| Donor 2 | 100 | 101.1 | 27.4 | 34.9 |
| Donor 3 | 100 | 116.0 | 20.7 | 29.6 |
| Donor 4 | 100 | 90.5 | 34.6 | 52.4 |
| Donor 5 | 100 | 144.6 | 84.1 | 102.6 |
| Donor 6 | 100 | 130.8 | 95.0 | 112.4 |
| Donor 7 | 100 | 137.5 | 110.3 | 85.8 |
| Donor 8 | 100 | 133.6 | 67.6 | 71.6 |
| Donor 9 | 100 | 172.7 | 51.3 | 85.0 |
| Donor 10 | 100 | 169.3 | 22.8 | 105.1 |

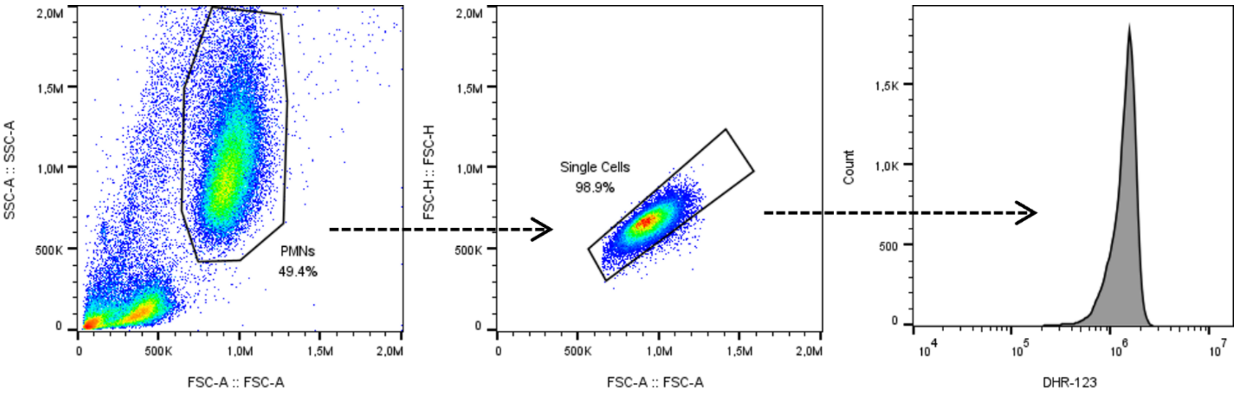


**Figure S1 – gating strategy for the DHR-123 experiments shown in figure 2.**

**Table S2 – cK58β2m and dK58β2m inhibit baseline ROS production in PMNs.** MFI values (x$10^4$) from individual donors from the DHR-123 experiment shown in figure 2.

| | Untreated | DPI | β2m | cK58β2m | dK58β2m |
|---|---|---|---|---|---|
| Donor 1 | 134.0 | 55.1 | 86.2 | 50.7 | 50.0 |
| Donor 2 | 116.0 | 56.7 | 128.0 | 71.9 | 77.6 |
| Donor 3 | 106.0 | 44.7 | 90.8 | 34.5 | 31.1 |
| Donor 4 | 55.9 | 32.5 | 52.4 | 33.4 | 37.3 |
| Donor 5 | 32.9 | 25.1 | 29.7 | 27.0 | 34.1 |
| Donor 6 | 55.7 | 25.3 | 63.6 | 24.9 | 32.0 |
| Donor 7 | 155.0 | 49.8 | 159.0 | 58.5 | 77.1 |
| Donor 8 | 66.3 | 26.3 | 59.3 | 31.0 | 51.4 |
| Donor 9 | 79.3 | 29.5 | 85.9 | 27.5 | 25.6 |
| Donor 10 | 18.5 | 14.6 | 20.2 | 16.2 | 22.4 |

**Table S3 – cK58β2m and dK58β2m inhibit fMLF-induced ROS production in PMNs.** MFI values (x$10^4$) from individual donors from the DHR-123 experiment shown in figure 2.

| | Untreated | fMLF | β2m + fMLF | cK58β2m + fMLF | dK58β2m + fMLF |
|---|---|---|---|---|---|
| Donor 1 | 61.9 | 123.0 | 113.0 | 100.0 | 100.0 |
| Donor 2 | 20.0 | 61.1 | 64.9 | 43.4 | 43.5 |
| Donor 3 | 12.9 | 61.6 | 71.9 | 28.5 | 26.2 |
| Donor 4 | 57.7 | 127.0 | 131.0 | 102.0 | 52.6 |
| Donor 5 | 26.0 | 85.9 | 107.0 | 85.6 | 79.0 |
| Donor 6 | 30.2 | 89.6 | 90.8 | 63.8 | 68.7 |
| Donor 7 | 50.5 | 110.0 | 107.0 | 94.1 | 59.9 |
| Donor 8 | 74.7 | 114.0 | 108.0 | 97.1 | 70.1 |
| Donor 9 | 67.3 | 106.0 | 110.0 | 90.2 | 78.4 |

**Table S4 – cK58β2m and dK58β2m inhibit TNF-α-induced ROS production in PMNs.** MFI values (x$10^4$) from individual donors from the DHR-123 experiment shown in figure 2.

| | Untreated | TNFα | β2m + TNFα | cK58β2m + TNFα | dK58β2m + TNFα |
|---|---|---|---|---|---|
| Donor 1 | 20.4 | 98.6 | 95.7 | 93.2 | 88.6 |
| Donor 2 | 10.2 | 76.0 | 74.6 | 50.3 | 23.8 |
| Donor 3 | 63.9 | 101.0 | 104.0 | 95.1 | 63.0 |
| Donor 4 | 21.9 | 103.0 | 102.0 | 78.1 | 73.5 |
| Donor 5 | 27.6 | 95.1 | 87.7 | 82.1 | 43.5 |
| Donor 6 | 46.6 | 113.0 | 109.0 | 109.0 | 93.4 |
| Donor 7 | 23.3 | 72.9 | 74.4 | 56.5 | 31.0 |
| Donor 8 | 16.2 | 78.4 | 68.9 | 47.5 | 12.8 |
| Donor 9 | 11.2 | 80.4 | 82.4 | 59.7 | 21.2 |
| Donor 10 | 29.6 | 101.0 | 109.0 | 95.1 | 60.6 |

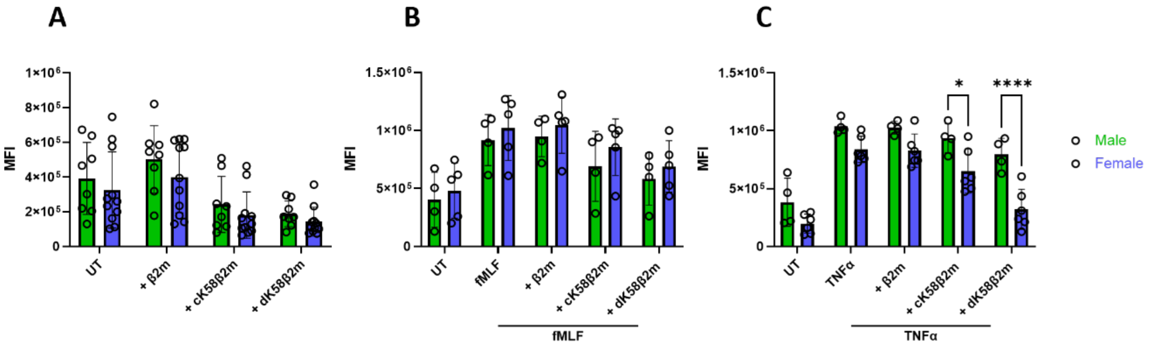


**Figure S2 – Differences in ROS production between male and female donors as assessed by DHR-123 fluorescence.** PMNs were treated with 50 µg/ml β2m, cK58β2m or dK58β2m either alone (A) or in combination with 1 µM fMLF (B) or 10 ng/ml TNF-α (C) for 1 h. Intercellular ROS were detected using DHR-123 (40 µM) by flow cytometry. Results are expressed as mean MFI ± SD. n=19 (A), n=10 (B), n=9 donors (C), *P<0.05, ***P<0.0001. Comparisons between male and female donors were analyzed with two-way ANOVA.

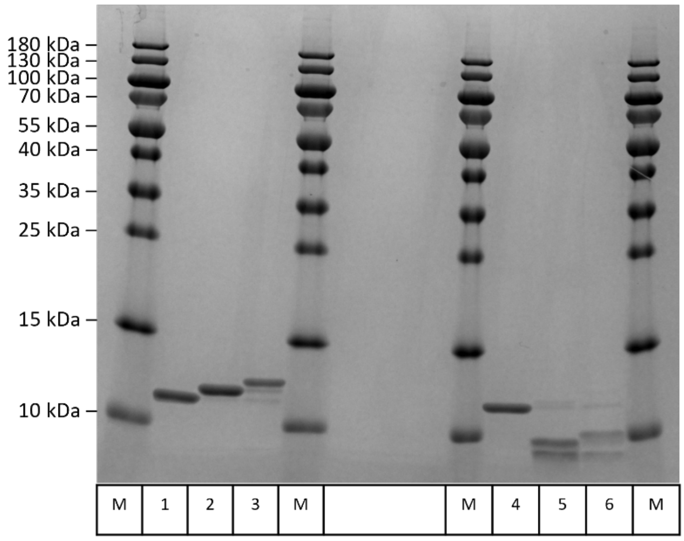


| | |
|---|---|
| M | Molecular marker |
| 1 | β2m non-reduced |
| 2 | cK58β2m non-reduced |
| 3 | dK58β2m non-reduced |
| 4 | β2m reduced |
| 5 | cK58β2m reduced |
| 6 | dK58β2m reduced |

**Figure S3 – Analysis of β2m, cK58β2m and dK58β2m under non-reduced and reduced conditions.** Under reducing conditions, cK58β2m and dK58β2m separates into two distinct chains, consistent with cleavage occurring within the disulfide loop of β2m. Trace amounts of native β2m can be seen present in cK58β2m and dK58β2m samples.
